# PISA: versatile microscope for 3D single molecule light sheet imaging

**DOI:** 10.1101/2024.12.05.625331

**Authors:** Yuichi Taniguchi, Kazuya Nishimura, Yamato Yoshida, Sooyeon Kim, Latiefa Kamarulzaman, David G. Priest, Masae Ohno

## Abstract

Light sheet microscopy combines high sensitivity and imaging speeds with low photodamage, however a remaining hurdle to its widespread adoption is its requirement for special sample preparation. Our new microscope - planar illumination microscope for single molecule imaging for all purpose (PISA) - significantly relieves these restrictions to enable light sheet imaging of coverslip-mounted samples as commonly analyzed in biological microscopes. We show how this technology now permits high-throughput imaging using multi-well plates. Further, the ability of 3D single molecule imaging in microlitre or mm^3^ scale volumes over the coverslip improves the sensitivity of biochemical gel-based fluorometric assays to atto/zepto molar levels. PISA will facilitate widespread adoption of light sheet imaging for a variety of biological, biochemical and medical investigations.

## Introduction

A grand challenge in biology is to explore the 3D distributions of the molecules that are the fundamental units of cells, and thereby build a bottom-up understanding of biological systems. Single molecule fluorescence imaging is a method that allows direct exploration of such molecular spatial distributions^1,2^ and has a wide range of applications. For example, *in vivo* it has been used to map 3D cellular structures such as of the mitochondria^3,4^, count molecular numbers for quantifying transcriptomes and proteomes^5,6^, and track molecular dynamics during protein complex assembly^7^. *In vitro* it is also used to characterise individual molecular dynamics, and for real-time DNA sequencing^8^.

The two main problems of single molecule imaging are the limited imageable sample volume and complex sample preparation. Conventionally, single molecule sensitivity was achieved by use of an evanescent field^2^ or highly inclined illumination microscopy^9^. However, with these techniques the imageable sample depth was limited to below a sub-micron or several micron above the coverslip. This largely restricts the analysis to proteins bound to a coverslip, or the underside of cells cultured on a dish. This depth can be extended to the sub-millimeter level using wide-field illumination microscopy, however the high out-of-focus fluorescence background limits observations to samples containing only very sparse fluorescence molecules at any moment. Under such conditions, the PALM/STORM technique can be used to build images by observing randomly activated fluorescence signals over a longer time^3,4^, however this hampers live-cell analysis and fast imaging. Recently, high depth and low background imaging has been achieved with light sheet microscopy^10^ coupled to high numerical aperture (NA) fluorescence detection^11–15^. Here however, usability is severely restricted because structural constraints require the sample to be encapsulated in special chambers, such as an agarose cylinder^10^, glass box^11^, or groove^13^ or in a large open dish that internalizes the objective lenses^12,14^ or a small mirror^15^.

Light-sheet microscopy has the significant unique advantage of being able to provide 3D sectioned images at higher speeds with lower phototoxicity and photobleaching^10–22^ as well as single molecule sensitivity. This outstanding performance suggests that once the structural limitations have been overcome, this microscopy could displace widely-used biological microscopes, such as a confocal and wide-field. Here, we present a microscope setup, called the <u>p</u>lanar <u>i</u>llumination microscope for single molecule imaging for <u>a</u>ll purpose (PISA), that overcomes the inherent obstacles of light-sheet microscopy to enable 3D single molecule imaging in general scientific research.

## Results

### Open-top light sheet microscope that allows 3D single molecule imaging

To approach the problem of a generalisable light-sheet microscope, we aimed for a similar user experience to conventional inverted biological microscopes whilst retaining the capability for single molecule observation using light sheet illumination. To realize this, we designed a microscope structure that enables measurement of common coverslip-mounted samples with no special procedures. First, we positioned the sample plane entirely above the optical systems for light sheet imaging, where two objective lenses for fluorescence illumination and detection are orthogonally placed below the coverslip position at a tilted angle (Fig. 1a and b). For high NA imaging, water immersion was used for both illumination and detection objective lenses by setting a custom-designed reservoir between them. To minimize optics misalignments due to thermal drift, the two objective lenses were physically fixed and conjugated on a custom-designed microscope body, and water or immersion medium was continuously supplied via the reservoir during longer time courses (Supplementary Fig. 2). We adopted a 20× objective lens (XLUMPLFLN 20XW, NA = 1.0, Olympus, Japan) to enable observation of a relatively large field of view while maintaining the sensitivity to detect single molecules. The sample holder was designed to minimize physical interference of the immersion reservoir thereby enabling efficient access to the whole coverslip area. In this setup, like conventional inverted microscopes, the imaging can be done simply by placing the sample on the stage above the objective lenses (Supplementary Fig. 3). The sample holder was placed on positional stages to translate the sample in 3D along either the detection or horizontal axes. Further, a transillumination system with a halogen lamp and condenser was placed above the sample along the detection axis.

**Figure 1.**
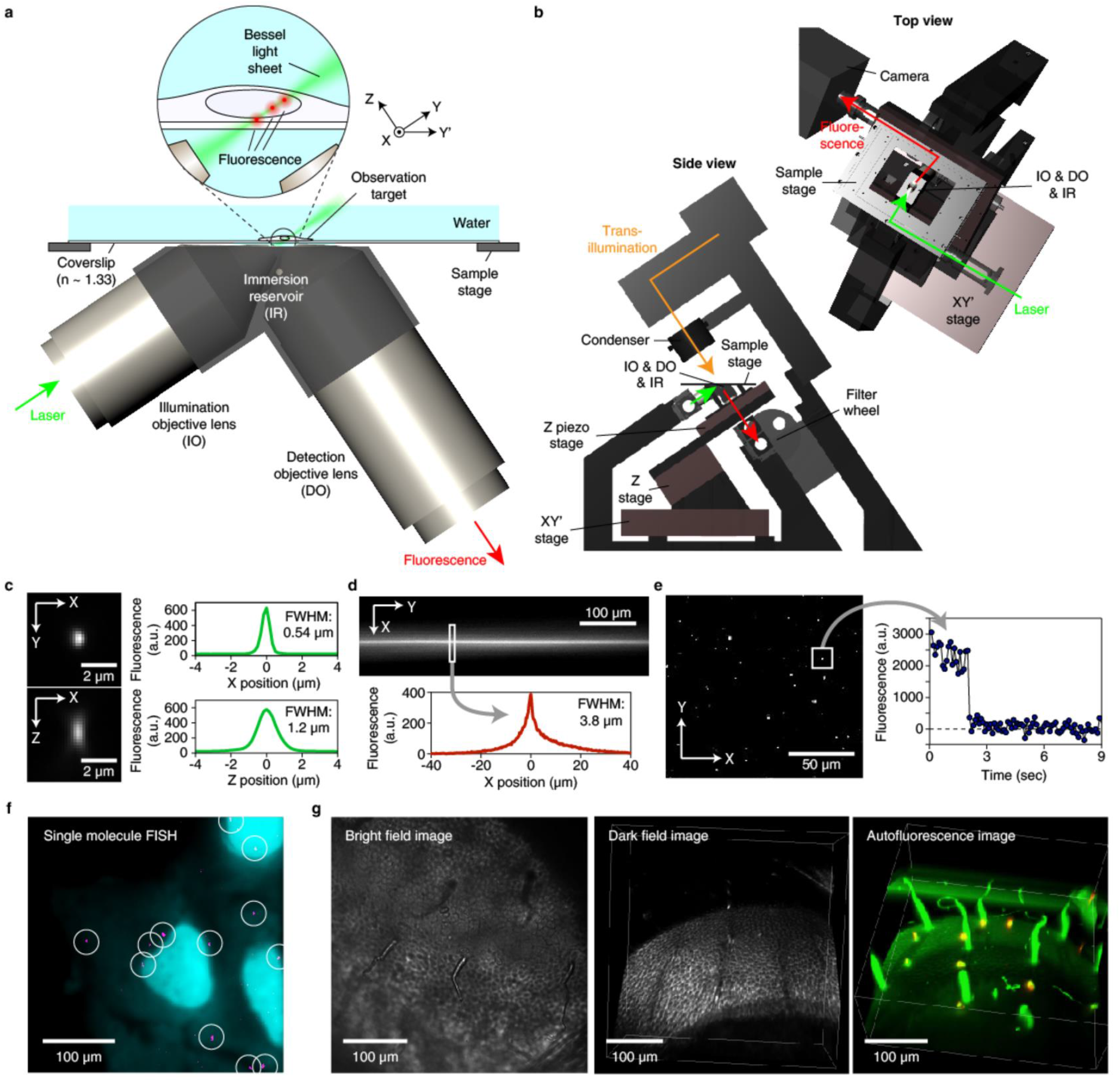
Overview of the PISA microscope. **a**, An expanded view of the microscope around the objective lens for fluorescence illumination and detection. The Y and Z axis is defined to be along the optical axis of the illumination and detection objective lens respectively. The X axis is defined perpendicularly against the Y and Z axis, while the Y’ axis is along the horizontal axis and is perpendicular to the X axis. The sample can be translated along the X, Y’ or Z axis, whilst the objective lenses are immobile. **b**, Side (left) and top (right) view of the microscope. **c**, A point spread function in the X-Y (top) and X-Z plane (bottom), obtained by imaging 0.04 μm fluorescent microbeads. Images (left) and fluorescence distributions at the center pixel of the point spread functions (right) are shown. **d**, An image (top) and fluorescence distribution along the X-axis (bottom) of the bessel beam light sheet. **e**, Visualization of single Atto 647 dye molecules fused in a gellan gum gel. One step photobleaching of an observed fluorescence spot is shown on the right. **f**, Visualization of *RDH14* RNA molecules in U2OS cells with single molecule FISH. **g**, Bright field (left), dark field (middle) and autofluorescence (right) observation of an *Oryzias latipes* egg.

An outstanding problem with this setup is that asymmetric optical aberration can occur for both illumination and detection pathways passing through the tilted coverslip region due to mismatched refractive indices between the coverslip and water. To eliminate this, we posited to use a special coverslip with a refractive index close to that of water (*n* = 1.33) and as thin as possible. Such a coverslip in single or multiple well format is commercially available as Lumox (SARSTEDT, Germany) or fluorocarbon film (Zell-kontakt GmbH, Germany), which were originally made for creating a gas permeable cell culture condition. We found that optical aberration was successfully eliminated enough to observe symmetric point spread functions through these films (Fig. 1c, Supplementary Fig. 4). The inherent autofluorescence of these films could be significantly reduced by irradiating with a UV light overnight.

We further integrated microscopy components to realize 3D single molecule imaging over as large a volume as possible. For this, a bessel beam light sheet^14,16^, constructed by scanning a focused laser with an axicon lens, was used for fluorescence illumination, yielding an even fluorescence excitation field over the entire field of view of the detection objective lens. And for fluorescence detection at the single molecule level we used an EM-CCD or sCMOS camera (Supplementary Fig. 1). We confirmed that our configuration with the special coverslip allows creation of a bessel beam with a 3.8 μm FWHM for ∼500 μm length (Fig. 1d). Using this setup, we successfully imaged single Atto 647 dye molecules in a gellan gum gel up to 0.2 mm depth over the coverslip (Fig. 1e). In this setup, 3D imaging can be done by stacking images acquired during movement of the sample along either the detection or horizontal axes using the positioning stages, which we call Z- and Y’-stacking, respectively. We confirmed that our system could take 3D Y’-stack images of single RNA molecules (single molecule FISH) in human cultured cells (Fig. 1f). In addition, the high sensitivity of the microscope coupled with stage scanning allowed us to take clear autofluorescence 3D images of *Oryzias latipes* (Japanese medaka killifish) eggs (Fig. 1g, right, Supplementary Movie 1). We also note that our setup further allows 3D-sectioned dark-field observation by detecting scattered light irradiated by the sheet illumination, as well as bright-field observation using the transillumination system (Fig. 1g, left and middle, Supplementary Movie 2).

### High-speed, high-sensitivity live cell 3D imaging in conventional biological samples

Live cell 3D imaging with low phototoxicity is an important advantage of the light sheet microscope, where laser irradiation is selectively restricted to the observed plane^10,17,18^. The PISA microscope achieves even lower phototoxicity due to its high fluorescence sensitivity, which allows more frequent image acquisition per unit time. We tested this ability by carrying out high-speed 3D time lapse imaging of embryogenesis in *Caenorhabditis elegans* eggs isolated from adult worms (Fig. 2a and b). By performing 3D time-lapse imaging at a rate of 30 frames per minute, we observed an exponential separation of chromosomes during cell divisions (Fig. 2c, Supplementary Movie 3). This is consistent with them being driven by elastic pulling forces from the poles. We also note that this experiment requires promptness because *C. elegans* eggs commence embryogenesis quickly after extraction from dissected adults (Fig. 2a). Many light sheet microscopes require samples to be first fused in an agarose gel, or, would require the floating egg positions to be perturbed by the objective lens or mirrors, making them unsuitable for time-sensitive experiments such as this. In contrast, our setup allows quick and efficient experiments equivalent to those performed using conventional inverted microscopes but at single molecule resolution (Fig. 2a).

**Figure 2.**
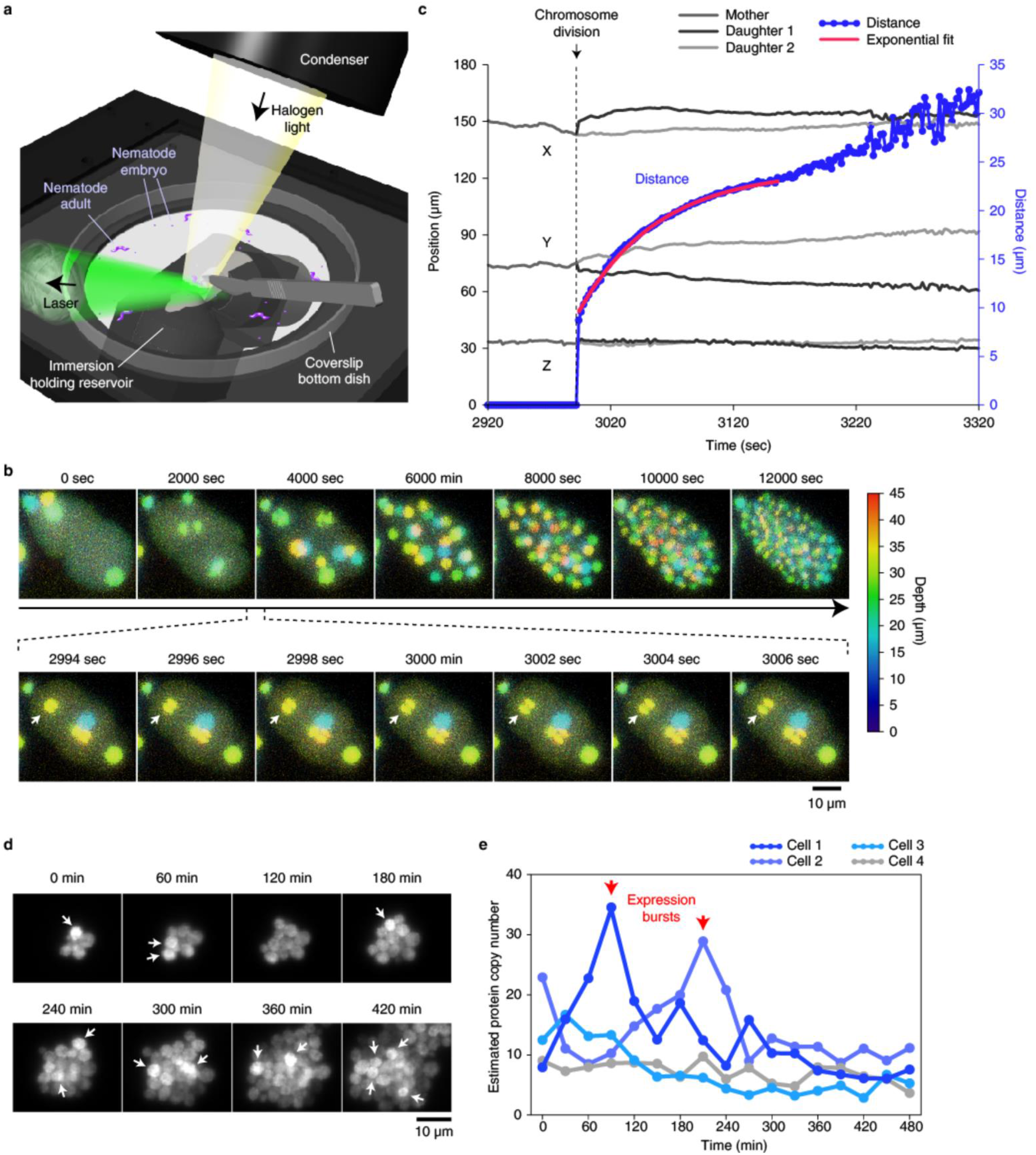
Low-phototoxicity, high-speed and high-sensitivity live cell 3D imaging with PISA. **a**, The open top design allowing direct handling and manipulation of measured samples as in conventional biological microscopes. **b**, High-speed 3D time lapse imaging of *Caenorhabditis elegans* embryogenesis. mCherry fused to histone H2B protein HIS-58 is imaged. Fluorescence signals at different depths are shown in different colors. **c**, Time course of 3D positions for the two divided chromosomes that are indicated in Fig. 2b with an arrow, and the distance between them. The time (*t*)-dependent change in the distance (*d*) can be described by an exponential equation (the red curve): *d* = 24.3 -14.6×*exp*(−0.031×(*t* - 2994)). **d**, Time-lapse imaging of expression of DDR48 protein molecules fused to Venus in single *Saccharomyces cerevisiae* cells. Bursty, pulsing expressions of proteins are indicated with arrows. **e**, Time course of estimated protein copy numbers in single cells. The copy numbers are estimated by dividing fluorescence counts with a value from single molecules, which was obtained by measuring a strain sparsely expressing membrane-bound Venus proteins.

Live cell imaging with the PISA microscope also provides an opportunity to discover and characterize dynamics of small numbers of molecules in cells. To demonstrate this, we imaged expression of DDR48 proteins genetically fused to fluorescent proteins in live *Saccharomyces cerevisiae* cells (Fig. 2d). The measurement was made possible by sandwiching cells between the coverslip and an agarose gel pad containing SC medium as done in standard yeast experiments, which is difficult in some types of light sheet microscopes. Only newly expressed proteins during the time-lapse intervals are imaged by exposing lasers a couple of times to cause photo-bleaching of existing fluorophores after every image acquisition. Imaging at 2 frames per hour, we observed bursty, pulsing protein expressions in single cells at different timings (Fig. 2d). By normalizing fluorescence signals with single molecule fluorescence counts, protein numbers in bursts were estimated to be 10-20 molecules (Fig. 2e). Note that because sheet illumination prevents photobleaching at non-imaged planes, quantitative 3D fluorescence signals can be characterized even under a strong laser irradiation to detect small numbers of fluorescent molecules across the entire cell volume.

### High-throughput light sheet imaging for multi-well plates over large volumes

Another important advantage of our setup is the ability to scan unconstrained volumes above the coverslip out to the extent of the working distance of the objective lens. This allows high-throughput imaging using multi-well plates or large-area scanning measurement, which has been challenging with existing light sheet microscopes. To demonstrate this, we performed high-throughput, high-content imaging of HeLa cells in multi-well plates treated with 96 different anti-cancer drugs from a library (SCADS inhibitor Kit 3.1, Screening Committee of Anticancer Drugs, Japan) (Fig. 3a). We obtained clear 3D sectioned images of fluorescently labeled nuclear membrane and genomic DNA in cells in each well (Fig. 3b), revealing a list of drugs that cause similar morphological changes (Fig. 3c, Supplementary Table 1). Further, we imaged a large number of HeLa cells in 3D by large-area Y’-stack imaging utilizing the long scanning distance of PISA (Fig. 3d). Note that, in contrast to conventional microscopes such as wide-field and confocal, Y’-stack imaging, with the tilted observation plane, allows large-area, stream acquisition scanning of the sample in 3D by continuous stage movement and camera acquisition, enabling fast cell population analysis analogous to flow cytometry. Considering that our setup can image a tilted plane of ∼500×500 μm size covering the 200 μm sample depth at each stack, we can analyze a 0.12 mm^3^ (= 0.5×1.2×0.2) volume per minute when acquiring Y’-stack with 1 μm steps at a 50 msec exposure time. As a result, we could typically analyze 3D fluorescence images of >2,000 cells in 10 minutes using our setup (Fig. 3e). This number can be further improved by increasing the camera acquisition speed or Y’-stack imaging area, as long as clear fluorescence signals are obtained. Together, these abilities of PISA will open the door to sophisticated cytometric analyses with high 3D resolution, minimized photodamage, multiple samples, and single molecule sensitivity.

**Figure 3.**
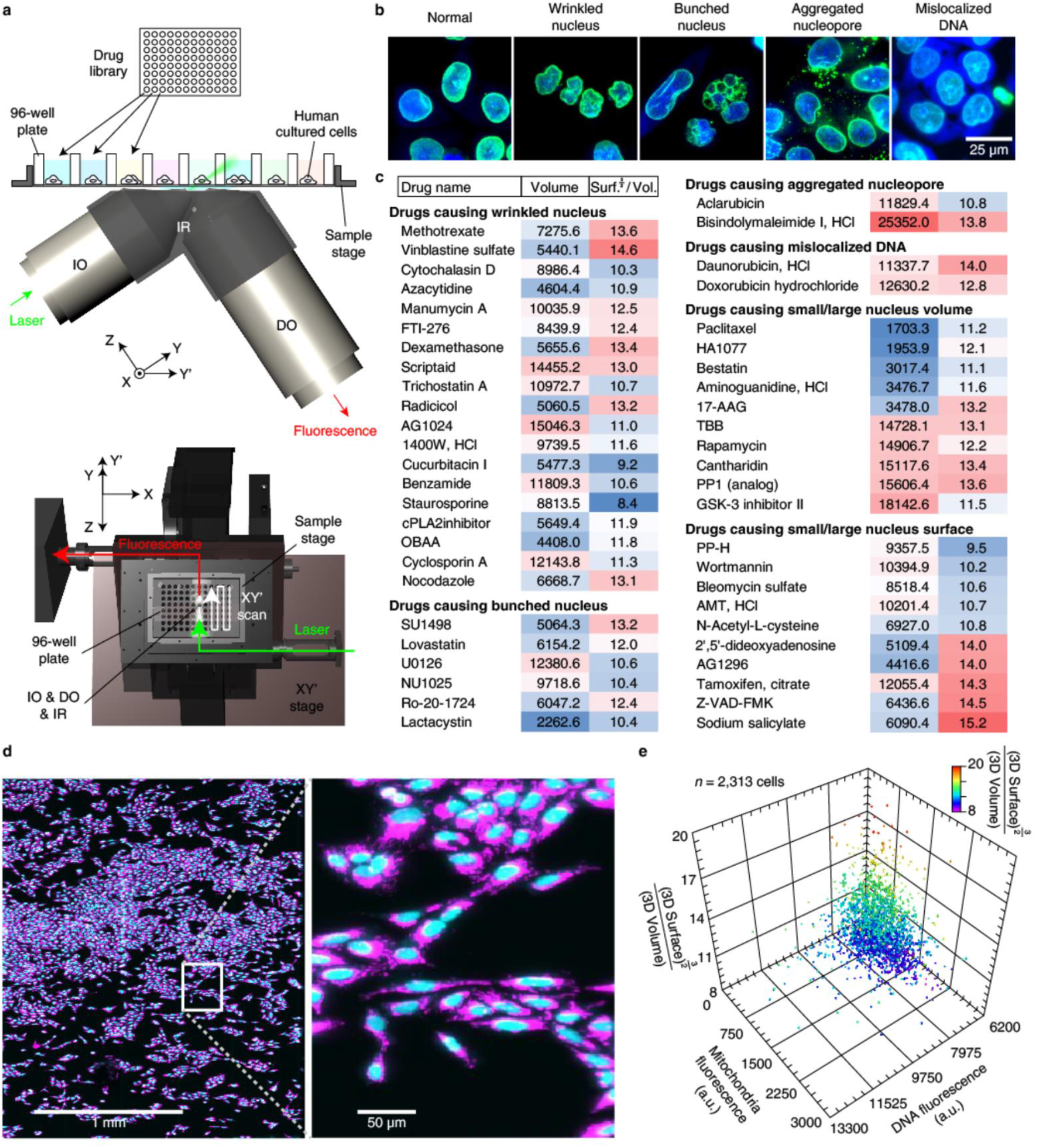
High-throughput and high-content imaging with PISA. **a**, The experimental system for high-content drug screening in HeLa cells. HeLa cells in a 96-well plate were treated with different anti-cancer drugs from a library for 3 hours and then imaged. **b**, Typical examples of cellular morphological changes with drug treatments. DNA stained with Hoechest (blue) and a nucleopore protein Pom121 fused to Venus (green) were imaged. Data from NL-71-101 (Normal), Vinblastine sulfate (Wrinkled nucleus), Ro-20-1724 (Bunched nucleus), Aclarubicin (Aggregated nucleopore) and Doxorubicin hydrochloride (Mislocalized DNA) are shown. **c**, A list of drugs causing specific morphological changes of cells. 3D volume, and a ratio of 3D surface (to the power of 3/2) to 3D volume are shown on the right. **d**, Large-area 3D Y’-stack imaging for dish-cultured U2OS cells. DNA and mitochondria in cells were stained by DAPI (blue) and MitoTracker (pink), respectively. **e**. 3D projection of quantitative 3D characteristics in individual cells.

### Applicability for single molecule biochemistry

The ability of mm^3^ or μL order volume imaging together with single molecule sensitivity suggests the possibility of detecting ultra-low, atto-molar (aM) or zepto-molar (zM) concentrations of molecules in biochemical samples. Further, the μL-scale imaging volume will allow bulk-scale biochemical fluorometric analyses with such ultra sensitivity, including protein/DNA concentration fluorometer measurement, SDS-PAGE or DNA electrophoresis, and flow cytometry. To test this ability, we applied single molecule detection to SDS-PAGE analysis (Fig. 4a), where we measured a polyacrylamide gel containing different concentrations of fluorescently labeled protein (Fig. 4b).

**Figure 4.**
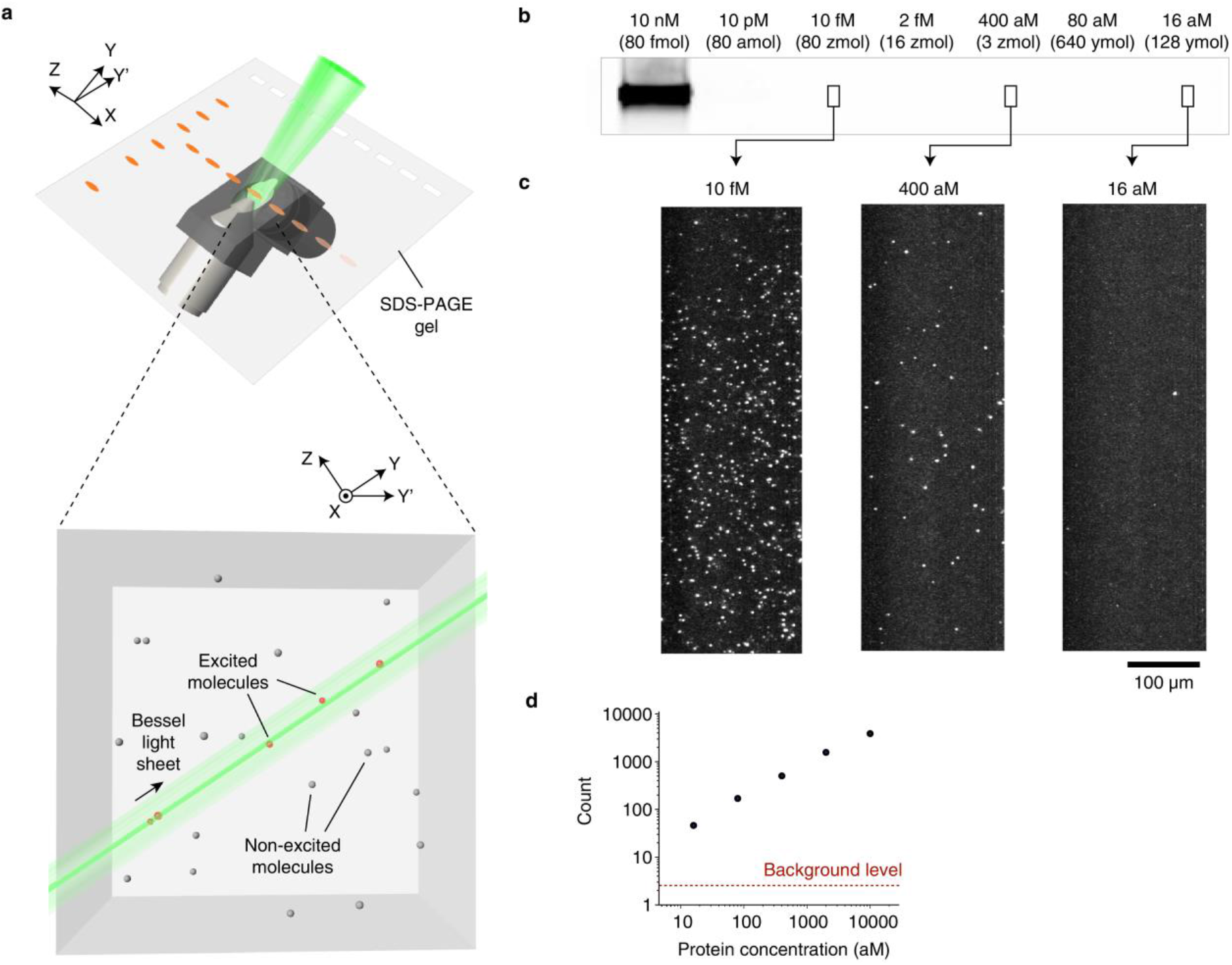
Single molecule detection in a biochemical assay with PISA. **a**, The experimental system for single molecule SDS-PAGE. **b**, An image of a polyacrylamide gel acquired with a standard gel imager (ImageQuant LAS 4000, GE healthcare). The gel lanes contain different concentrations of fluorescently-labeled protein from 10 nM (far left) to 16 aM (far right). **c**, Representative single molecule fluorescence images of the gel with PISA microscopy. Images were obtained by deskewing and z-projection of the Y’-stacks. **d**, Linearity of single molecule counts on protein concentrations. Background level was obtained from counts of empty gel images.

While a conventional gel imager detected a band up to 10 nM, our setup detected bands up to atto- or zepto-molar levels while maintaining signal linearity (Fig. 4c and 4d). This indicates that, in theory, ∼10^9^ times higher sensitivity can be achieved using our setup. We expect that this measurement can be extended to an ultra-sensitive Western blotting assay for detecting, for example, the presence of disease-causing proteins, and small number, or single cell proteome analysis. Furthermore, characterization of individual molecules will yield novel dimensions of parameters different from bulk methods such as the brightness or spectrum of individual labeled bio-molecules.

## Discussion

Taken together, our setup, PISA, realizes both the usability of biological inverted microscopes and the performance of single molecule light sheet microscopes. Since the optical pathway is simple, we can easily integrate advanced optical methods for light sheet imaging such as a lattice light sheet^12^, two photon illumination^16,19^, stimulated emission depletion^20^ and airy beam^21^, which may further improve the spatial resolution. Further, by obviating the need for special sample chambers, PISA can widely inherit practical advantages of conventional microscopes, such as measuring large-sized samples, imaging the coverslip surface, and prevention of dirt sticking to the objective lens. PISA will position the single molecule light sheet microscope for widespread adoption as a general tool in the biological, biochemical and medical research communities.

## Methods

### Microscope

The PISA microscope was built on a custom microscope body designed using 3D CAD software (Rhinoceros, Robert McNeel & Associates, USA) and crafted in a metalworking company (Iwamoto Model Co., Japan) (Supplementary Fig. 1). The microscope body has two components. The first component is a base to mechanically connect the two water-immersion objective lenses for fluorescence illumination (54-10-7, NA = 0.66, 28.6×, Special Optics, USA) and detection (XLUMPLFLN 20XW, NA = 1.0, 20×, Olympus, Japan) tilted at 33.8 degrees on a vibration-free table (MS-4000, Newport, USA). This tight connection can be crucial, since the initial version of the PISA microscope (Supplementary Fig. 5) that can adjust the two objective lens positions independently using optics mounts suffered from thermal drift and vibration of the objective lenses. The two objective lenses are also connected via a custom-designed water immersion reservoir sealed with silicone (Dent silicone V, Shofu, Japan). For immersion, either water or immersion medium (Immersion medium Immersol W 2010, Zeiss, Germany) are used. An extruded part of the illumination objective lens on the immersion reservoir is shaved with a file so as not to interfere with the coverslip. The illumination port of the microscope is connected to laser sources as described later. The laser source is passed through an axicon lens (Mie Optics, Japan) to generate a bessel beam and is reflected with a galvano mirror (6215HB, Cambridge Technology, USA) to generate a light sheet. The detection port is connected to a sCMOS camera (ORCA-Flash4.0 V2, Hamamatsu Photonics, Japan), or an EM-CCD camera (iXon Ultra 897, Andor, UK) through an imaging lens. A motorized filter wheel (FW102C, Thorlabs, USA) with dichroic filters (Semrock, USA) is inserted in the front of the imaging lens. The second component is a multi-dimensional stage to move the sample along the horizontal and detection axis against the immobile objective lenses. For the horizontal movement, an XY motorized stage (H117, Prior Scientific Instruments, UK) was mounted on the vibration-free table. Meanwhile, for wide-range and rapid movements along the detection axis, a motorized (MLJ050/M, Thorlabs, USA) and piezoelectric (NanoScanZ, Prior Scientific Instruments, UK) stage, respectively, was mounted on the XY stage tilted at 15 degrees. A sample holder is set on the stage, and the holder can mount a gas chamber that can be connected to a mixed CO_2_ gas supply (TK-0003MIGM, Tokken, Inc., Japan). A custom-made incubator box combined with an air heater (TK-0003HU20, Tokken, Inc., Japan) is placed to cover the entire microscope system.

For the laser sources, we used 405 nm diode laser (MLL-III-405-100mW, Changchun New Industries Optoelectronics Technology, China), 488/514.5 nm Argon ion laser (Innova70-C, Coherent, USA), 570 nm fiber laser (2RU-VFL-P-1000-570, MPB Communications, Canada), 592 nm fiber laser (VFL-P-1000-592, MPB Communications) and 647 nm fiber laser (2RU-VFL-P-300-647, MPB Communications). These laser sources are combined with dichroic mirrors and were coupled with an optical fiber. The fiber-coupled laser is emitted on a breadboard that is set parallel to the illumination axis. The laser emission is switched by mechanical shutters (VMM-D3, Vincent Associates, USA).

Imaging software (MetaMorph, Molecular Devices, USA) and programs written in LabVIEW (National Instruments, USA) with a multifunction I/O device (USB-6363, National Instruments, USA) is used to synchronously control the camera, stages, filters, and shutters in order to carry out the experiments. Image analyses were performed using ImageJ.

For the coverslip, we used a commercial Lumox (SARSTEDT, Germany) or fluorocarbon (FC) film (Zell-kontakt GmbH, Germany) with a single or multiple well format. Autofluorescence of these films was eliminated by irradiating with UV overnight, or by coating a thin agarose gel on the coverslip. Alternatively, we used a lab-made Cytop (Asahi Glass Co., Japan) film, which also has a refractive index similar to water.

### Measurement of point spread functions and light sheet distributions

The point spread function was measured by imaging 0.04 μm fluorescent beads (FluoSpheres® Carboxylate-Modified Microsphere yellow-green fluorescent (505/515), Life Technolgies) embedded in 1% agarose gel by a 514.5 nm excitation through a filter (FF01-542/27-25, Semrock) (Fig. 1c). For each PSF, 121 Z-scanning series of images were acquired with 0.125 μm increment per plane, and the averaged intensity profiles were calculated from 5 time measurement with ImageJ software (NIH, USA). The bessel beam spatial distribution was measured by imaging solution of Atto 647 dye (Sigma-Aldrich) under a 647 nm excitation through a filter (FF01-708/75-25, Semrock) (Fig. 1d).

### Single molecule FISH

A U2OS cell line was cultured on a 24-well FC plate (Zell-kontakt GmbH, Germany) in FluoroBrite Dulbecco’s modified Eagle’s medium (DMEM) (Thermo Fisher Scientific) supplemented with 10% fetal bovine serum (FBS) and 1× GlutaMAX™-I (GIBCO) at 37°C with 5% CO_2_ for 48 to 72 hours. The cultured cells were fixed in 4% paraformaldehyde in 1× phosphate buffered saline (PBS) for 15 min and were washed with 1× PBS and methanol. The fixed cells were immediately transferred to a −20°C freezer until use. The stocked cells were briefly washed once with ice-cold 1× PBS, were permeabilized for 5 min with 0.25% (v/v) Triton X-100 in 1× PBS at room temperature, and were washed three times with ice-cold 1× PBS. The cells were incubated for 5 min in wash buffer containing 2× saline-sodium citrate buffer (SSC) (Sigma) and 20% formamide (Nakalai). Cells were then hybridized with the primary oligonucleotide probes that contain complementary sequences against *RDH14* RNA and the secondary probes. The probes were designed using the probe design tool Stellaris® probe designer (https://www.biosearchtech.com/stellaris-designer). The hybridization was done with 200 μl of 10 nM primary probes in hybridization buffer containing 2× SSC, 20% formamide, 0.2 mg/ml bovine serum albumin, 100 mg/ml dextran sulfate (Sigma), 2 mM vanadyl ribonucleoside complex (Sigma), and 1 mg/ml *Escherichia coli* tRNA (Sigma) in a humid chamber inside a 37°C-hybridization oven for 12 hours. Following hybridization, the cells were washed twice with wash buffer every 30 minutes followed by two washes with 2× SSC at room temperature. The washed cells were postfixed in 4% paraformaldehyde in 2× SSC at room temperature for 30 min. The fixed cells were washed three times with 2× SSC and once with wash buffer. The cells were then hybridized with the secondary probes that are labeled with Cy5 at the 5’ ends and are complementary to the primary probes. The hybridization was performed by gently adding 200 μl of 2 nM secondary probes in hybridization buffer. The cells were incubated in a humid chamber inside a 37°C-hybridization oven for 12 hours. The cells were washed twice with wash buffer every 30 minutes, followed by two washes with 2× SSC at room temperature. Cells were then imaged with the PISA microscope under a 647-nm laser illumination through a filter (FF01-708/75-25, Semrock) (Fig. 1e).

### Observation of embryogenesis in *Caenorhabditis elegans*

*Caenorhabditis elegans* strain oxSi487 [mex-5p::mCherry::H2B::tbb-2 3’UTR::gpd-2 operon::GFP::H2B::cye-1 3’UTR + unc-119(+)] II (Caenorhabditis Genetics Center, EG6787, ref. 23) was raised at 20°C on a nematode growth medium agar plate (1.7% (w/v) agar, 50 mM NaCl, 0.25% (w/v) peptone, 1 mM CaCl_2_, 5 μg/mL cholesterol, 25 mM KH_2_PO_4_, 1 mM MgSO_4_) seeded with *Escherichia coli* OP50, and were retrieved into M9 medium. Bodies of single worms were cut with a knife to take out embryos. Embryos from each worm were separately transferred into each well in a FC plate, and were imaged with PISA microscope (Fig. 2c). 3D time-lapse imaging was carried out every 2 minutes for 7.5 hours at RT. Each Z-stack 3D image was acquired with 1 μm step size for total 70 μm thick with 10 msec exposure per frame. Imaging was performed with a 570 nm laser excitation via a filter (FF01-617/73, Semrock, USA).

### Observation of protein expression dynamics in *Saccharomyces cerevisiae*

The *Saccharomyces cerevisiae* strain we imaged was constructed from a BY4741 strain by replacing genomic DDR48 coding sequence with 4×Venus sequence. Cells were cultured in Synthetic complete (SC) medium at 30°C overnight, and were washed with SC medium. Cells were then placed on a 50-mm Lumox dish (SARSTEDT, Germany) and were sandwiched with an agarose gel pad containing SC medium. Images were acquired every 30 minutes at RT (Fig. 2d). Each Z-stack 3D image was acquired with 0.5 μm step size for total 40 μm thick with 10 msec exposure per frame. Imaging was performed with a 514.5 nm laser excitation via a filter (FF01-542/27-25, Semrock).

To obtain an estimated protein copy number, we divided observed fluorescence counts by a value from a single molecule. The single molecule value was obtained by imaging a strain expressing Venus-PIL1 fusion proteins under an uninduced Gal1 promoter, which causes membrane-localized, sparse single molecule spots on the cell membrane, and by analyzing an averaged fluorescence count of the single spots.

### High-content drug screening in HeLa cells

A HeLa cell line expressing Pom121-Venus was cultured in DMEM (D5796, Sigma Aldrich) with 10 % FBS (Biowest, France) under 5% CO_2_ at 37°C. Cultured cells were suspended in 250 μL of low fluorescence DMEM (FluoroBrite™ DMEM, Thermo Fisher Scientific) to a final concentration of 10^5^ cells/mL, and 100 μL suspension was pipetted into each well in a 96-well FC plate (Zell-kontakt GmbH, Germany), and were cultured for 24 hours at 37°C. Each 10 μL of the anti-cancer drug library (SCADS inhibitor Kit 3.1, Japan) was added to each well to a final concentration of 10 μM, and the plate was incubated for 3 hours at 37°C. Cells were then fixed with 3.7 % formaldehyde at RT for 20 min. After two times washes with PBS, 50 μL of 1 μg/mL Hoechst 33258 (Dojindo, Japan) in PBS was added to each well and was incubated at RT for 30 min, after which cells were washed twice with PBS. Pom121-Venus was excited with 514.5 nm laser, and was detected through a band-pass filter (545BP40, Omega Optical) (Fig. 3b). Hoechst 33258 was imaged with 405 nm laser through a band-pass filter (FF01-483/32-25, Semrock).

### Large-area imaging in U2OS cells

A U2OS cell line was cultured on a 24-well FC plate (Zell-kontakt GmbH, Germany) in FluoroBrite DMEM (Thermo Fisher Scientific) supplemented with 10% FBS and 1× GlutaMAX™-I (GIBCO) and incubated at 37°C with 5% CO_2_ for 48 to 72 hours. The cultured cells were fixed in 4% paraformaldehyde in PBS for 15 min and washed with PBS. The fixed cells were then stained with DAPI and MitoTracker (Thermo Fisher Scientific). Y’-stack 3D image was acquired with 1 μm step size with 30 msec exposure per frame (Fig. 3d). DAPI and MitoTracker was imaged with a 405 and 647 nm laser excitation via a filter (FF01-464/542/639-25 and FF01-708/75-25, Semrock), respectively.

### Single molecule SDS-PAGE

A polyacrylamide gel for SDS-PAGE was created by following the standard protocol, and was irradiated with UV overnight. Electrophoresis was performed in the polyacrylamide gel loading different amounts (80 fmol to 128 ymol) of a SeTau647-labelled transferrin (Fujifilm Wako, Japan) in SDS sample buffer, prepared with a serial dilution. The gel was first scanned with ImageQuant LAS4000 (GE Healthcare) (Fig. 4b). The gel was then imaged with the PISA microscope in Y’-stack with 3 μm step size at 50 msec exposure per frame (Fig. 4c). Imaging was performed with a 647 nm laser excitation via a filter (FF01-708/75-25, Semrock).

## Data availability

The datasets generated during and/or analyzed during the current study are available from the corresponding author on reasonable request.

## Acknowledgments

The authors thank Y. Okada, Y. Saito and T. Masujima for critical advices to the microscope; K. Kyoda, J. Takayama and S. Onami for assisting *C. elegans* experiments; L. Simon and V. Kumar for helpful discussions; M. Ishida for experimental assistance; and H. Pickersgill from Life Science Editors for editorial assistance. HeLa cell line which expresses Pom121-Venus were provided from N. Imamoto in RIKEN, Advanced Science Institute. *Oryzias latipes* eggs were provided from N. Sakata in RIKEN, Quantitative Biology Center. Anti-cancer drug library was provided from Screening Committee of Anticancer Drugs, Japan. This work was supported by grants-in-aid for Scientific Research (A) (20H00460) and Young Scientists (A) (24687022), Challenging Pioneering Research (19H05545 and 20K20458) and Exploratory Research (26650055), Early-Career Scientists (19K15718 and 22K14800) and Scientific Research on Innovative Areas (23115005 and 20H05936); Japan Society for the Promotion of Science; and CREST (JPMJCR2334), PRESTO (JPMJPR15F7) and ACT-X (JPMJAX1914); Japan Science and Technology Agency; and grants from the RIKEN DECODE project, Stage Transition project, RIKEN Incentive Research Projects; the Takeda Science Foundation; Suntory Rising Stars Encouragement Program in Life Sciences (SunRiSE).

## Contributions

Y.T. designed the microscope. Y.T. and K.N. constructed the microscope system. Y.T., K.N., Y.Y., S.K. and L.K. performed imaging experiments. M.O. contributed to cell preparations and data managements. Y.T. and D.P. wrote the paper.

